# The versatile regulation of K_2P_ channels by polyanionic lipids of the phosphoinositide (PIP_2_) and fatty acid metabolism (LC-CoA)

**DOI:** 10.1101/2021.07.01.450694

**Authors:** Elena B. Riel, Björn C. Jürs, Sönke Cordeiro, Marianne Musinszki, Marcus Schewe, Thomas Baukrowitz

**Affiliations:** Institute of Physiology, Christian-Albrechts University of Kiel, 24118 Kiel, Germany; MSH Medical School Hamburg, University of Applied Sciences and Medical University, 20457 Hamburg, Germany

**Keywords:** K_2P_ channel, KCNK, K_2P_ channel regulation, lipid gating, polyanionic cellular lipids, long-chain fatty acid ester (oleoyl-CoA), PI(4,5)P_2_ (PIP_2_)

## Abstract

Work of the past three decades provided tremendous insight into the regulation of K^+^ channels - in particular K_ir_ channels - by polyanionic lipids of the phosphoinositide (e.g. PIP_2_) and fatty acid metabolism (e.g. oleoyl-CoA). However, comparatively little is known regarding the phosphoinositide regulation in the K_2P_ channel family and the effects of long-chain fatty acid CoA esters (LC-CoA, e.g. oleoyl-CoA) are so far unexplored. By screening most mammalian K_2P_ channels (12 in total), we report strong effects of polyanionic lipids (activation and inhibition) for all tested K_2P_ channels. In most cases the effects of PIP_2_ and oleoyl-CoA were similar causing either activation or inhibition depending on the respective subgroup. Activation was observed for members of the TREK, TALK and THIK subfamily with the strongest activation by PIP_2_ seen for TRAAK (~110-fold) and by oleoyl-CoA for TALK-2 (~90-fold). In contrast, inhibition was observed for members of the TASK and TRESK subfamilies up to ~85 %. In TASK-2 channels our results indicated an activatory as well as an inhibitory PIP_2_ site with different affinities. Finally, we provided evidence that PIP_2_ inhibition in TASK-1 and TASK-3 channels is mediated by closure of the recently identified lower X-gate as critical mutations within the gate (i.e. L244A, R245A) prevent PIP_2_ induced inhibition. Our results disclosed K_2P_ channels as a family of ion channels highly sensitive to polyanionic lipids (PIP_2_ and LC-CoA), extended our knowledge on the mechanisms of lipid regulation and implicate the metabolisms of these lipids as possible effector pathways to regulate K_2P_ channel activity.

## Introduction

Members of the large family of two-pore domain potassium (K_2P_) channels are critically involved in many cellular functions ranging from renal ion homeostasis, cell development, hormone secretion, immune functions as well as cardiac and neuronal excitability (Enyedi and Czirjak, 2010). Accordingly, dysregulation of K_2P_ channels is seen in many disease states such as cardiac disorders (i.e. atrial fibrillation, ventricular tachycardia) (Liang et al., 2014; Decher et al., 2017b), hyperaldosteronism (Davies et al., 2008; Bandulik et al., 2015), pulmonary arterial hypertension (Olschewski et al., 2006; Ma et al., 2013; Antigny et al., 2016), as well as pain perception disorders such as migraine and depression (Alloui et al., 2006; Heurteaux et al., 2006; Lafreniere et al., 2010; Andres-Enguix et al., 2012; Royal et al., 2019). Initially thought to mediate passive background conductance in excitable cells, increasing evidence shows that K_2P_ channels are highly regulated by sensing a broad range of diverse physiological stimuli and endogenous ligands. However, not all K_2P_ channels respond to the same set of stimuli but rather their sensitivity profile corresponds to their affiliation to one of the known six subfamilies (TREK, TASK, TALK, THIK, TWIK and TRESK). Members of the TREK subfamily (TREK-1, TREK-2 and TRAAK) display the most diverse (and e.g. regarding TREK-1 the best investigated) regulation with relevant stimuli including temperature (Maingret et al., 2000a), mechanical force (Maingret et al., 1999; Chemin et al., 2005), membrane voltage (Maingret et al., 2002; Schewe et al., 2016), extracellular/intracellular pH (pHe/pHi) (Maingret et al., 1999; Honore et al., 2002), partner proteins (Plant et al., 2005; Sandoz et al., 2006) and various lipids (Maingret et al., 2000b; Chemin et al., 2005; Chemin et al., 2007). Members of the TALK subfamily (TASK-2, TALK-1 and TALK-2) are activated by high pHe (Morton et al., 2003; Duprat et al., 2005; Niemeyer et al., 2010). TASK-1, TASK-3 and TASK-5 make up the TASK subfamily and members are inhibited by extracellular acidification (Rajan et al., 2000; Bayliss et al., 2001; Morton et al., 2003) but also respond to membrane lipids such as diacylglycerol (DAG) (Wilke et al., 2014). Members of the TWIK subfamily (TWIK-1, TWIK-2 and TWIK-3) are rather stimuli insensitive but their cellular activity is controlled by regulated protein trafficking (Feliciangeli et al., 2007; Feliciangeli et al., 2010) as well as possibly SUMOylation (Rajan et al., 2005; Plant et al., 2010) and unique features of SF gate (Nematian-Ardestani et al., 2019). The TRESK subfamily has just one member (i.e. TRESK) and is regulated in particular by changes in intracellular Ca^2+^ levels mediated by Calmodulin-dependent phosphatase calcineurin (Czirjak et al., 2004). For the THIK subfamily (THIK-1 and THIK-2), no physiological gating stimulus has been reported so far.

Work of the last three decades revealed that many ion channels are regulated by phosphoinositides and in particular by the most abundant phosphoinositide phosphoinositol-4,5-bisphosphate (PI(4,5)P_2_, PIP_2_). PIP_2_ sensitive channels include all members of the inward rectifying K_ir_ channels (Suh and Hille, 2005; Logothetis et al., 2007; Suh and Hille, 2008) but also members of the voltage gated K_v_ channels (Oliver et al., 2004; Rodriguez et al., 2010; Kruse et al., 2012; Zaydman and Cui, 2014; Taylor and Sanders, 2017), transient receptor potential TRP cation channels (Qin, 2007; Suh and Hille, 2008), Ca^2+^-activated BK-type channels (Vaithianathan et al., 2008) and a number of ion transporters (i.e. NCX, NCE, PMCA) (Hilgemann et al., 2001; Gamper and Shapiro, 2007). Accordingly, the significance of PIP_2_ regulation of ion channels is demonstrated by physiologically important processes such as insulin secretion in pancreatic β-cells (Rorsman and Ashcroft, 2018) or mechanical transduction and adaptation in hair cells (Hirono et al., 2004; Effertz et al., 2017). The mechanisms of regulation in some of these channels have been resolved to the atomic level (Hansen et al., 2011; Niu et al., 2020; Sun and MacKinnon, 2020).

The first report on the regulation of a K_2P_ channel by phosphoinositides dates to 2005 and still represents a main reference on this topic (Chemin et al., 2005). Chemin and colleagues reported the strong activation of TREK-1 channels by several phospholipids including PE, PI, PC and PIP_2_ and identified a cluster of basic residues in the proximal C-terminus as potential PIP_2_ binding region (Chemin et al., 2005). Accordingly, it was proposed that negatively charged lipids (e.g. PIP_2_) electrostatically attract the C-terminus towards the membrane and, thereby, cause channel activation as also confirmed later using a fluorescently labelled C-terminus (Sandoz et al., 2011). Later the same authors reported that PIP_2_ application can also produce TREK-1 inhibition but the reason for this variability remains unresolved (Chemin et al., 2007). More recently, an additional PIP_2_ interaction region in a more distal C-terminus of TREK-1 channels was identified (Soussia et al., 2018). Aside the TREK subfamily, the effects of PIP_2_ have also been studied in TASK-1, TASK-2, TASK-3 and TRESK channels. Initially, the inhibition of TASK-1 and TASK-3 by PLC activation was thought to result from PIP_2_ break down (Lopes et al., 2005) as PIP_2_ appeared to cause activation, however, later it became clear that DAG (released by PLC) likely inhibit TASK-1 and TASK-3 channels directly (Wilke et al., 2014). In TASK-2 channels and human (but not rod) TRESK channels application of PIP_2_ to excised patches has been reported to cause activation (Lopes et al., 2005; Niemeyer et al., 2017; Giblin et al., 2018).

Another class of polyanionic lipids known to regulate ion channels are long-chain fatty acid Coenzym esters (LC-CoA). These obligate metabolites of cellular fatty acids affect many K_ir_ channels producing activation in K_ATP_ (Kir6.2/SUR) channels (Branstrom et al., 1998; Schulze et al., 2003) but inhibition in most other K_ir_ channels (Shumilina et al., 2006; Tucker and Baukrowitz, 2008). Aside the K_ir_ channel family, little is known about the effects of LC-CoA on other ion channels. However, TRPV1 channels have been reported to be activated by LC-CoA (Yu et al., 2014). In both, K_ir_ channels and TRPV1, LC-CoA appear to interact with the same sites that also bind PIP_2_ likely via the negative phosphate groups present in both types of lipids (Schulze et al., 2003; Yu et al., 2014).

Aside the aforementioned exception of TREK/TRAAK, TASK-1/-2/-3 and TRESK channels, effects of phosphoinositides on other members of the K_2P_ channel family have not been investigated yet, and K_2P_ channel regulation by LC-CoA is so far unexplored. Therefore, we systematically studied the effect of PIP_2_ and LC-CoA on K_2P_ channels by testing all functionally expressing channels (12 out of 15) under identical conditions and quantified their respective sensitivities. We uncovered that all K_2P_ channel members strongly respond to at least one of the two polyanionic lipid species and that the six K_2P_ channel subfamilies can be classified as either polyanionic lipid-activated or lipid-inhibited subfamilies. Further, we investigated the physicochemical prerequisites for lipid activation in TALK-2 channels and the structural regulation mechanism of lipid regulation in the TASK subfamily. We finally discuss the potential physiological relevance of our findings that, however, warrant further investigation in more native preparations.

## Results

### PIP_2_ causes subtype-dependent responses (activation/inhibition) in most K_2P_ channels

We studied the direct effect of the most abundant phosphoinositide PI(4,5)P_2_ (PIP_2_) on all known functionally expressing K_2P_ channels (12 in total) by applying 10 μM PIP_2_ to the cytoplasmatic site of the respective channels in inside-out patches excised from *Xenopus* oocytes. The type of response (activation or inhibition) varied depending on the K_2P_ channel subfamily and in the efficacy of activation or inhibition depended on the particular subfamily member (**Figure 1A**). Robust PIP_2_ activation was observed within the TREK subfamily with 17 ± 3-fold, 11 ± 4-fold and 114 ± 24-fold activation for TREK-1, TREK-2 and TRAAK, respectively (**Figure 1 A**). The PIP_2_-activated currents displayed slightly outward rectifying current-voltage (I-V) relationships between −80 mV and +40 mV (**Supplementary Figure S1A-C**).

**Figure 1.**
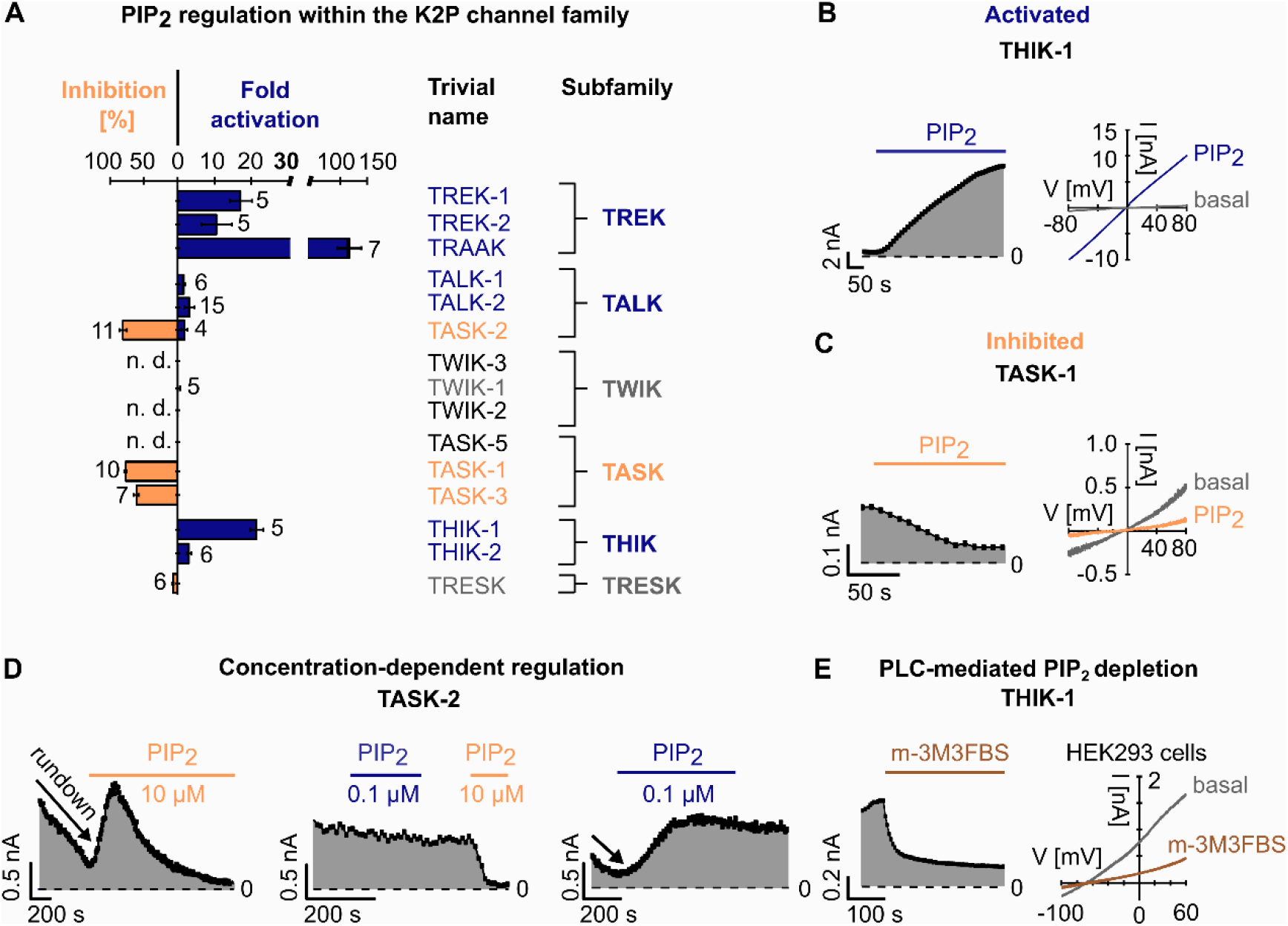
PIP_2_ regulation of K_2P_ channels. **A.** Fold activation (blue) and inhibition [%] (orange) of K_2P_ channel currents by 10 μM PIP_2_ at +40 mV, measured in ramp protocols as shown in B and C. Insensitive channels are highlighted in grey. n. d. = not determined. Number (n) of independent experiments is indicated next to the bars. **B,C.** Representative current traces (right) and analysed currents at +40 mV plotted over time (left) of THIK-1 (B) and TASK-1 (C) channels measured in voltage ramps between −80 and +80 mV in excised inside-out patches of *Xenopus* oocytes using control bath solution and in the presence of 10 μM PIP_2_. **D.** Analysed TASK-2 currents at +40 mV plotted over time, measured as in B,C, using control bath solution and in the presence of 0.1 μM or 10 μM PIP_2_. Current rundown is indicated with an arrow. Low PIP_2_ concentration (0.1 μM) rescues current rundown (right) but produces no inhibition (middle); 10 μM PIP_2_ also rescue rundown if present (left) but leads to a subsequent inhibition of the channel. If no rundown is present 10 μM PIP_2_ only lead to inhibition (middle). **E.** Representative traces of THIK-1 currents in whole-cell experiments using HEK293 cells measured in voltage ramps between - 100 and +60 mV, analysed at 0 mV. Measurements are performed in control solution and in the presence of 20 μM of the PLC activator m-3M3FBS. All data is presented as mean ± s.e.m.

Members of the THIK subfamily were also activated by PIP_2_ with strong activation (19 ± 2-fold) for THIK-1 and weaker activation (4 ± 1-fold) for THIK-2 and, the activated currents showed approximately linear I-V relationships (**Figure 1A,B, Supplementary Figure S1H**). Given the large PIP_2_ activation of THIK-1 we tested whether PIP_2_ break-down would cause current inhibition in a cellular context. Indeed, direct pharmacological activation of phospholipase C (PLC) using m-3M3FBS caused strong inhibition of THIK-1 currents measured in HEK293 cells (**Figure 1E**).

In the TALK subfamily PIP_2_ caused a 4 ± 1-fold activation of TALK-2 and a weaker activation (1.7 ± 0.2-fold) of TALK-1 currents (**Figure 1A, Supplementary Figure S1D,E**). The effect on TASK-2 was more complex and depended on the experimental history. In many excised patches TASK-2 currents showed strong current rundown and the application of 10 μM PIP_2_ produced initially a 2.4 ± 0.3-fold activation. However, this activation ceased with time and channel activity finally dropped below the starting level resulting in an effective PIP_2_ inhibition of 83 ± 3 % (**Figure 1A,D left panel**). In membrane patches lacking current rundown, PIP_2_ application (10 μM) caused inhibition without the initially activation (**Figure 1D middle panel**). Further, lower PIP_2_ concentrations (e.g. 0.1 μM) prevented current rundown (**Figure 1D middle panel**) and activated rundown currents without producing subsequent inhibition (**Figure 1D right panel**). These findings indicate two distinct regulatory sites in TASK-2 with lower PIP_2_ levels supporting basal channel activity via an activatory site and higher PIP_2_ levels causing inhibition possibly via a distinct inhibitory site.

In the TWIK subfamily, only TWIK-1 expressed functionally but lacked a PIP_2_ response (**Figure 1A, Supplementary Figure S1F**).

Likewise, no significant PIP_2_ response was observed in TRESK channels (the only member of this K_2P_ channel subfamily) (**Figure 1A, Supplementary Figure S1I**) in contrast to a previous report (Giblin et al., 2018).

Finally, in the TASK subfamily only TASK-1 and TASK-3 expressed functionally (TASK-5 is non-functional, (Ashmole et al., 2001)) and both channels showed marked inhibition (82 ± 2 % and 61 ± 4 %) upon PIP_2_ application (**Figure 1A,C, Supplementary Figure S1G**). In some patches TASK-1 channels also showed current rundown but in contrast to TASK-2, PIP_2_ failed to reactivate these channels (data not shown).

In summary, 10 of the 12 K_2P_ channels investigated changed activity upon PIP_2_ application with only TWIK-1 and TRESK lacking a significant PIP_2_ effect. Members of the TREK, THIK and TALK subfamilies were activated by PIP_2_ (TASK-2 only initially), whereas members of the TASK subfamily were inhibited.

### Most K_2P_ channels are highly sensitive to the fatty acid metabolite oleoyl-CoA

We further explored the K_2P_ channel lipid sensitivity by testing the polyanionic lipid oleoyl-CoA, which represents a common cellular long-chain fatty-acid metabolite. It is known to regulate ion channels (Rapedius et al., 2005; Ventura et al., 2005; Shumilina et al., 2006), however, it has not been investigated in K_2P_ channels so far. Upon application of 10 μM oleoyl-CoA strong activation of TREK-1 (15 ± 3-fold) and weaker activation of TREK-2 (6 ± 1-fold) was observed (**Figure 2A, Supplementary Figure S2A,B**). Notably, as seen with PIP_2_ (**Figure 1A, Supplementary Figure S1C**), TRAAK showed the highest lipid sensitivity within its subgroup, with oleoyl-CoA causing a 40 ± 16-fold current increase (**Figure 2A,B**). Further, the these activations were readily reversed upon extraction of oleoyl-CoA via the fatty acid binding protein BSA (**Figure 2B, Supplementary Figure S2A,B**).

**Figure 2.**
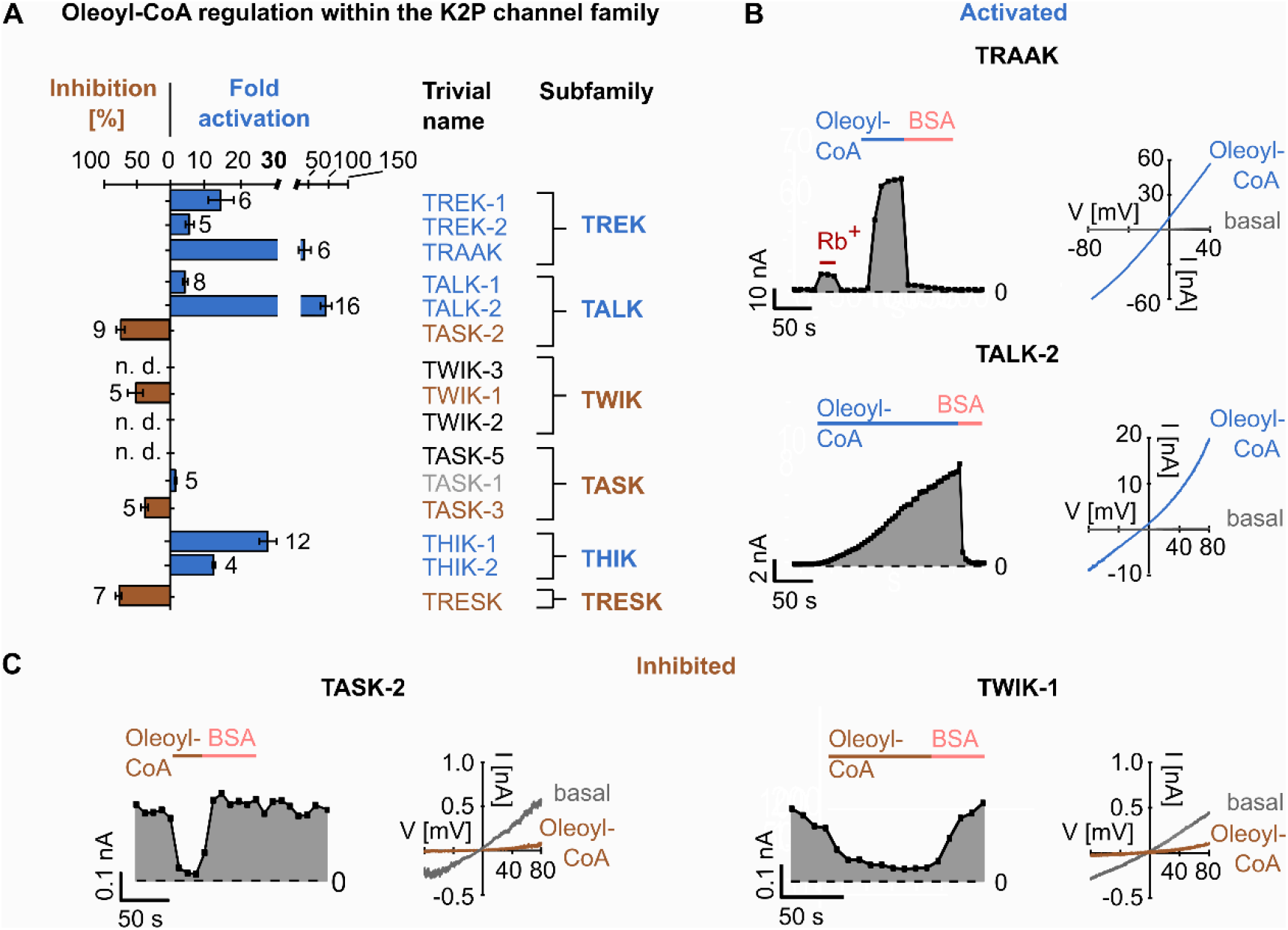
Oleoyl-CoA regulation of K_2P_ channels. **A.** Fold activation (blue) and inhibition [%] (brown) of K_2P_ channels at +40 mV by 10 μM oleoyl-CoA measured as in B and C. Insensitive channels are highlighted in grey. n. d. = not determined. Number (n) of independent experiments is indicated next to the bars. **B.** Representative current traces (right) and analysed currents at +40 mV plotted over time (left) of TRAAK and TALK-2 channels measured in voltage ramps between −80 and +80 mV in excised inside-out patches of *Xenopus* oocytes using control bath solution and in the presence of 10 μM oleoyl-CoA or 5 mg*ml^−1^ bovine serum albumin (BSA). **C.** Representative current traces of TASK-2 and TWIK-1 channels measured and analysed as in B. All data is presented as mean ± s.e.m.

Similar to the PIP_2_ responses, THIK-1 channels were strongly activated by oleoyl-CoA (29 ± 2-fold), whereas activation in THIK-2 was weaker (13.0 ± 0.4-fold) (**Figure 2A, Supplementary Figure S2F,G**) and thus, similar to the PIP_2_ responses.

In the TALK subfamily, TALK-2 stood out as oleoyl-CoA produced massive activation (94 ± 14-fold) while TALK-1 was only moderately activated (5 ± 1-fold) (**Figure 2A,B, Supplementary Figure S2C**). In contrast, TASK-2 channels were inhibited by oleoyl-CoA (70 ± 7 %) (**Figure 2A,C**), again similar to the PIP_2_ effects. Notably, in contrast to PIP_2_ (**Figure 1D**), oleoyl-CoA did not reactivate TASK-2 channels after current rundown (data not shown) suggesting that the potential activatory site in TASK-2 is selective for PIP_2_, whereas the potential inhibitory site appears to bind both types of lipids.

In the TASK subfamily, TASK-3 channels were inhibited by oleoyl-CoA (39 ± 5 %) in contrast to TASK-1 channels that were rather slightly activated (**Figure 2A, Supplementary Figure S2D,E**).

In the TWIK and TRESK subfamilies, both expressing members (i.e. TWIK-1 and TRESK) were markedly (~50 to ~75 %) inhibited by oleoyl-CoA (**Figure 2A,C, Supplementary Figure S2H**). As seen with all oleoyl-CoA effects reported here application of BSA reversed the action of oleoyl-CoA.

In summary, with the exception of TASK-1, oleoyl-CoA application modulated the activity of all K_2P_ channels tested. Further, the subfamily specific response (i.e. activation vs. inhibition) resembled to that seen with PIP_2_ in most cases. Exceptions to this rule were seen (i) for TASK-1 which was inhibited by PIP_2_ but not by oleoyl-CoA as well as (ii) for TWIK-1 and TRESK which were inhibited by oleoyl-CoA but lacked PIP_2_ sensitivity.

### Physico-chemical requirements of LC-CoA activation in TALK-2 channels

Given the particularly high sensitivity of TALK-2 to oleoyl-CoA (**Figure 3A**) we chose this channel to explore the LC-CoA properties required for activation in more detail. We obtained a dose-response relationship for oleoyl-CoA that was fitted to a standard Hill equation with an EC50 of ~11 μM (**Figure 2B**, **Supplementary Figure S3A-D**). Remarkably, however, even concentrations as low as 100 nM already produced robust (> 3-fold) TALK-2 channel activation (**Supplementary Figure S3A**) consistent with the high maximal effect (i.e. > 90-fold activation) at saturating concentrations (**Supplementary Figure S3C,D**). Further, the potency to activate TALK-2 channels negatively correlated with the number of LC-CoA double bonds present in fatty acids of 18 carbon atoms chain length, with the saturated stearic acid being the most potent LC-CoA (**Figure 3B,D**). Moreover, for saturated fatty acids, strongest activation was observed for stearyl-CoA, with palmityl-CoA and docosanoyl-CoA were less potent (**Figure 3C,D**). These results (except the activity drop for docosanoyl-CoA) can be explained by the lipid/water partitioning coefficient, as increasing acyl-chain length and reducing the number of double bonds is expected to promote membrane incorporation and, thus, would result in higher effective concentrations in the membrane from where channel interactions are likely to occur. The drop of potency for docosanoyl-CoA (compared to stearyl-CoA) apparently conflicts with this concept, suggesting that specific interactions of the fatty acid chain with the TALK-2 channel might also be important and, accordingly, may be less optimal for docosanoyl-CoA compared to stearyl-CoA.

**Figure 3.**
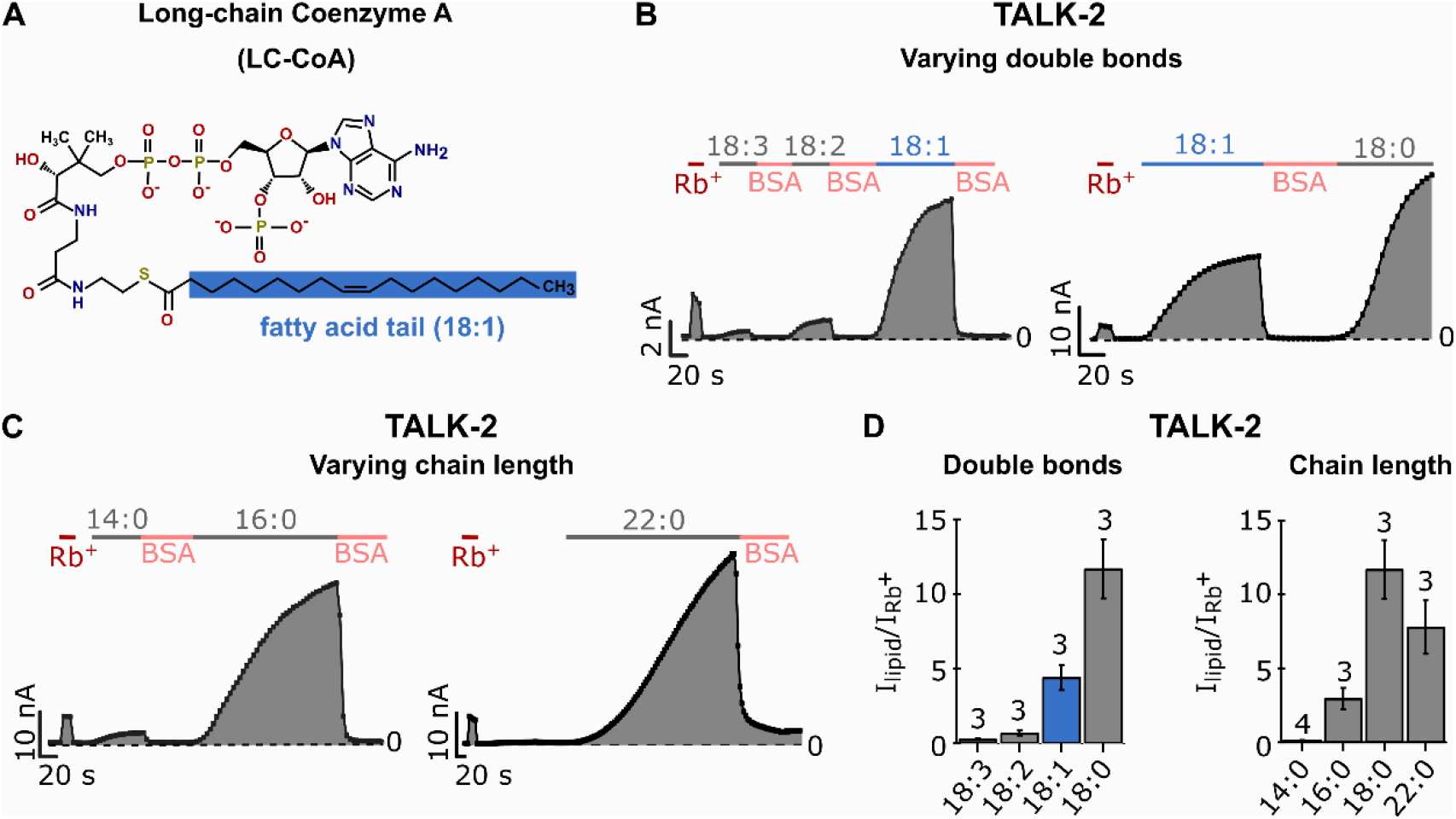
Physico-chemical properties of LC-CoA esters required for activation of TALK-2 K_2P_ channels. **A.** Chemical structure of long-chain Coenzyme A (oleoyl-CoA; 18:1), the fatty acid tail is shaded blue. **B.** Analysed current traces at +80 mV of TALK-2 channels measured in voltage ramps between −80 and +80 mV in excised inside-out patches of *Xenopus* oocytes using control bath solution and in the presence of 3 μM of long-chain fatty CoA esters with varying number and position of double bonds (18:0, 18:1, 18:2, 18:3) in the fatty acid tail of the molecule or 5 mg*ml^−1^ BSA. **C.** Analysed current traces at +80 mV of TALK-2 channels, measured as in B, using control bath solution an in the presence of 3 μM long-chain fatty acid CoA esters with varying chain length (14:0, 16:0, 18:0, 22:0) of the fatty acid tail. **D.** Fold activation of TALK-2 channels by long-chain fatty acid CoA esters with varying double bonds or chain length, measured as in B and C and normalised to the respective Rb^+^-activated current (red, B and C) at +80 mV. The average fold activation by Rb^+^ (I_Rb_^+^/I_basal_) of TALK-2 WT channels is 12 ± 3 (n = 20). Number (n) of independent experiments is indicated above the bars. All data is presented as mean ± s.e.m.

### Location of the PIP_2_ inhibition gate in TASK-1 and TASK-3

Here we report the inhibition of TASK-2, TASK-1, TASK-3, TWIK-1 and TRESK by the polyanionic lipids PIP_2_ and oleoyl-CoA. This raises the question of the nature of the inhibition gate. Interestingly, TASK-1 and TASK-3 have been previously shown to be inhibited by diacylglycerol (DAG) and a region named the halothane response element (HRE) in the distal TM4 segment was identified to be critical (Wilke et al., 2014). Moreover, in TASK-1 a gate at the HRE has recently been crystallographically identified and named the X-gate (Rodstrom et al., 2020). This X-gate forms a permeation blocking pore constriction at the contact points of the TM4 helices, located halfway between the selectivity filter (SF) gate and the cytoplasmic pore entrance. Mutations within the lower X-gate (e.g. L244A and R245A) (**Figure 4D**) have been shown to disturb channel closure resulting in channels with a higher relative open probability (Rodstrom et al., 2020). Notably, we found that the mutations L244A and R245A completely abolished PIP_2_ inhibition in TASK-1 and TASK-3 channels (**Figure 4A-C**) suggesting that PIP_2_ induces closure of the X-gate similar to DAG.

**Figure 4.**
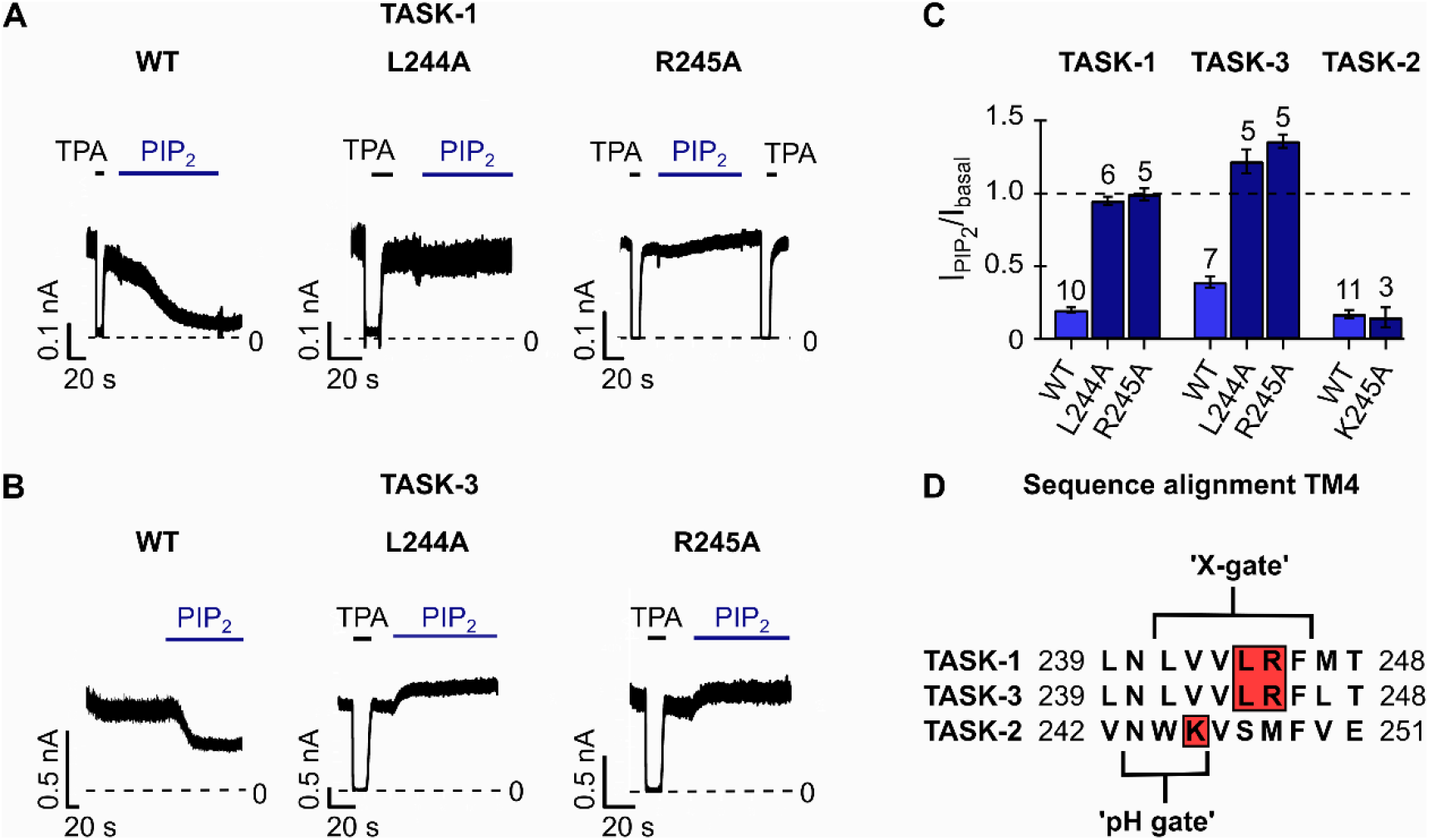
Mutations within the lower gate (X-gate) prevent PIP_2_ inhibition in TASK-1 and TASK-3 K_2P_ channels. **A,B.** Representative current traces of (A) TASK-1 WT, L244A and R245A mutant channels and (B) TASK-3 L244A and R245A mutant channels measured at continuous +40 mV in excised inside-out patches of *Xenopus* oocytes using control bath solution and in the presence of 10 μM PIP_2_ or 1 mM TPA, showing that mutations disrupting the lower gate render the channels insensitive to PIP_2_. **C.** Fold change of TASK-1, TASK-3 and TASK-2 WT (light blue) and mutant (dark blue) currents analysed at +40 mV in the presence of 10 μM PIP_2_ measured as in A and B, Figure 1D middle panel or Supplementary Figure S1J. Number (n) of independent experiments is indicated above the bars. **D.** Sequence alignment of TM4 of TASK-1, TASK-3 and TASK-2 K_2P_ channels. The lower X-gate in TASK-1 (TASK-3) and the region that forms the lower ‘pH gate’ in TASK-2 are highlighted. Red boxes show the location of introduced mutations in the respective K_2P_ channel. All data is presented as mean ± s.e.m.

In the pH sensitive TASK-2 channel a lower gate at a location similar to the X-gate in TASK-1 has been identified recently in cryo-EM structures obtained at different pH (Li et al., 2020). This work identified a lysine residue (i.e. K245) within the lower gate as a pH sensing residue and also as critical component of the lower gate itself. However, mutation of this lysine to alanine (K245A) had no obvious effect on PIP_2_ inhibition in TASK-2 (**Figure 4C,D, Supplementary Figure S2J**). This suggest that PIP_2_ inhibition is either mediated via a different gate (e.g. the SF) or that K245 is not critical for mediating PIP_2_ inhibition via the lower gate in contrast to pH inhibition.

## Discussion

Our comprehensive screen established K_2P_ channels as a family of K^+^ channels highly sensitive to polyanionic membrane lipids such as PIP_2_ and oleoyl-CoA. Further, depending on the particular K_2P_ subfamily polyanionic lipids either produced activation (TREK, TALK and THIK subfamily) or inhibition (TASK, TWIK and TRESK subfamily). The responses evoked in a particular subfamily produced by PIP_2_ and oleoyl-CoA were similar, however, with three exceptions: (i) TWIK-1 and (ii) TRESK were PIP_2_ insensitive, but were inhibited by oleoyl-CoA (iii) TASK-1 was inhibited by PIP_2_ but oleoyl-CoA insensitive (**Figure 5B**).

**Figure 5.**
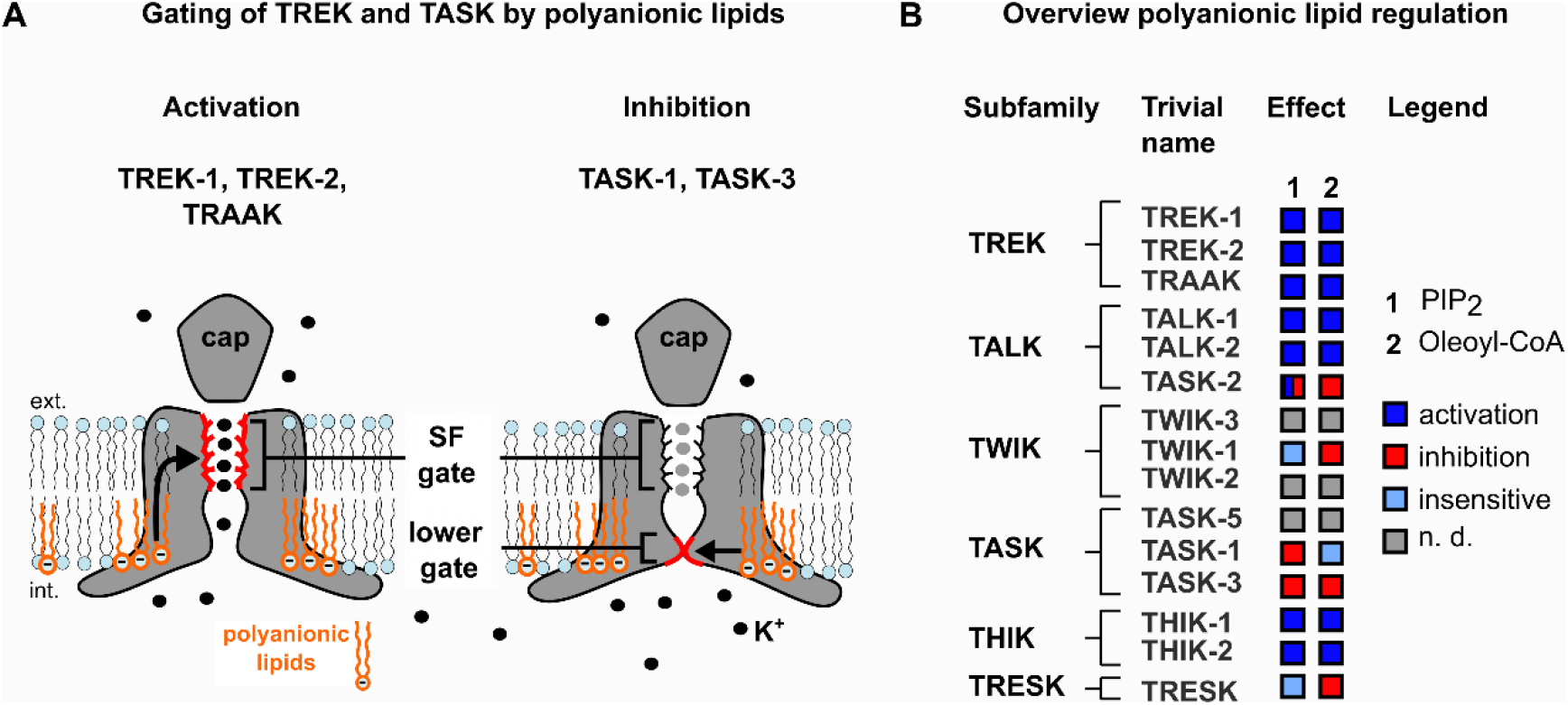
Regulation of K_2P_ channels by lipids. **A.** Cartoon illustrating the gating sites affected by polyanionic lipids (i.e. PIP_2_, oleoyl-CoA) in TREK/TRAAK and TASK channels. TREK channels adopt the active state at the selectivity filter (SF) in response to polyanionic lipids (orange). In contrast, TASK channels are inhibited by polyanionic lipids, that stabilise closure of the intracellular lower gate X-gate. **B.** Overview of the lipid regulation of K_2P_ channels: dark blue box = activation, red box = inhibition, light blue box = insensitivity to the lipids with (1) PIP_2_ and (2) oleoyl-CoA. The grey box indicates that the respective lipid effect was not determined (n. d.) caused by low functionally channel expression.

### Physiological implications of the PIP_2_ regulation in K_2P_ channels

In the K_ir_ channel family all members (i.e. K_ir_1.x, K_ir_2.x, K_ir_3.x, K_ir_4.x and K_ir_5.x) are thought to require PIP_2_ as mandatory co-factor to be functional (Huang et al., 1998; Logothetis et al., 2007; Furst et al., 2014). For K_2P_ channels not all members are activated by PIP_2_ and thus, PIP_2_ can be considered a modulatory agent rather than a mandatory co-factor. Accordingly, G protein coupled receptor (GPCR; i.e. P_2_Y_2_) activation of PLC produced strong inhibition of TREK-1 activity (i.e. activity is no more detectable) in HEK-293 cells, however, subsequent activating stimuli (e.g. temperature increase) still produced robust activation (**Supplementary Figure S1K**) indicating that PIP_2_ is not strictly required for channel activity.

The activation of TREK/TRAAK channels by PIP_2_ has been shown previously (Chemin et al., 2005; Chemin et al., 2007; Soussia et al., 2018). However, other studies report inhibition of TREK-1 channels by PIP_2_ (Cabanos et al., 2017) as well as split responses with PIP_2_ causing activation as well as inhibition (Chemin et al., 2007). In our hands inhibition of TREK-1/-2 channels or TRAAK channels was never observed in the *Xenopus* oocytes expression system. Moreover, we report here that TRAAK channels exhibited the strongest PIP_2_ response of all K_2P_ channels with a more than 110-fold increase in channel activity. The physiological relevance of this PIP_2_ regulation is currently unexplored but warrants further investigations in native preparations such as neurons which strongly express TRAAK channels (Brohawn et al., 2019; Kanda et al., 2019).

A notable finding of this work is the strong activation of THIK-1 channels by PIP_2_ (**Figure 1A,B**) as it represents the first reported physiological activator for these channels. Despite its expression in many regions of the CNS the specific role of THIK-1 channels is currently unknown with the exception of microglial cells, where THIK-1 activation was shown to be involved in microglial immune surveillance und inflammatory cytokine release (Madry et al., 2018). Interestingly, PIP_2_ production in microglia has been reported to play a key role in immune response signalling (Nguyen et al., 2017; Desale and Chinnathambi, 2021), and thus, possibly involves PIP_2_ activation of THIK-1.

We report here the inhibition of TASK-1 and TASK-3 channels by PIP_2_. The involvement of the PIP_2_ metabolism in the regulation of TASK-1 and TASK-3 has be intensively studied before and it is currently assumed GPCR mediated release of DAG directly inhibit these channels while the breakdown of PIP_2_ appears to be not critical (Bista et al., 2015). Thus, the activation of TASK-1 and TASK-3 via membrane PIP_2_ depletion (i.e. release of PIP_2_ inhibition by e.g. PLC activation) may not be of physiological relevance. However, inhibition of TASK-1 or TASK-3 via a local or global production of PIP_2_ or its enrichment in microdomains might be a regulatory mechanism.

TASK-2 channels have been previously reported to be activated by the short chain PIP_2_ derivative dioctanoyl-PIP_2_ (Niemeyer et al., 2017) which apparently contradicts to the here reported inhibition of TASK-2 by application of the native (i.e. long chain) PI(4,5)P_2_. However, we found that the PIP_2_ effect is clearly concentration-dependent with lower PIP_2_ concentrations supporting TASK-2 activity consistent with the reactivation of rundown of TASK-2 channel currents. Longer PIP_2_ applications as well as application on patches lacking current rundown produced robust and reliable inhibition. These findings indicate that distinct regulatory PIP_2_ sites exist in TASK-2 channels, i.e. a ‘higher’ affinity activatory PIP_2_ site and a ‘lower’ affinity inhibitory PIP_2_ site. Accordingly, at higher PIP_2_ concentration the inhibitory site dominates causing inhibition in TASK-2 (as seen with TASK-1 and TASK-3), whereas lower PIP_2_ concentration support TASK-2 activity via the higher affinity activatory site.

### Physiological implications of the LC-CoA regulation in K_2P_ channels

LC-CoA represent ubiquitous cellular products of the fatty acid metabolism as fatty acids bound to CoA before they can be taken up by mitochondria for β-oxidation. Our study revealed the regulation of many K_2P_ channels by oleoyl-CoA. LC-CoA has been previously reported to modulate several members of the K_ir_ channel family (Larsson et al., 1996; Rohacs et al., 2003; Rapedius et al., 2005; Shumilina et al., 2006; Tucker and Baukrowitz, 2008). Thereby, K_ATP_ channels are strongly activated while most other K_ir_ channels are inhibited likely because LC-CoA competes with PIP_2_ for binding, but lacks its activatory effect (competitive antagonism) (Shumilina et al., 2006). Such competitive antagonism, however, is unlikely to cause the here reported inhibition in TASK-2, TASK-1, TASK-3, TWIK and TRESK channels as these channels were not activated by PIP_2_. However, in the PIP_2_ activated channels (e.g. TREK-1) oleoyl-CoA likely interacts with the same sites as PIP_2_ because the degree of oleoyl-CoA and PIP_2_ activation is correlated in most K_2P_ channels.

The role of LC-CoA activation of K_ATP_ channels has been implicated in the mechanism of insulin secretion in pancreatic beta cells, in fatty acid sensing in hypothalamic neurons and in protection of cardiac myocytes under ischemic condition (Corkey et al., 2000; Liu et al., 2001; Tarasov et al., 2004; Le Foll et al., 2009; Rorsman and Ashcroft, 2018). These situations have in common that they cause an accumulation of LC-CoA in the cytoplasm which, potentially could also modulate the activity of the K_2P_ channels present in the respective tissues. In following a number of potential implications are pointed out that, however, will require further exploration in native preparations.

TALK-2 and TREK-1 channels are expressed in atrial as well as ventricular myocytes in the heart (Decher et al., 2001; Tan et al., 2004; Decher et al., 2017a). These channels are highly sensitive to LC-CoA as oleoyl-CoA caused a more then 15-fold activation (**Figure 2A,B, Supplementary Figure S1A,E**). Therefore, activation of TALK-2 and TREK-1 channels under ischemic conditions may have cardio protective effects in ventricular myocytes by shortening of action potential (AP) and concomitant reduction in Ca^2+^ loading (similar to KATP channels). However, this ischemic activation of TALK-2 and TREK-1 in the atrium causing AP shortening could potentially be also arrhythmogenic.

Noteworthy, TAKL-2 (together with TALK-1) is expressed in pancreatic β-cells, but its physiological role is currently unknown (Duprat et al., 2005; Rorsman and Ashcroft, 2018; Graff et al., 2021). Thus, the accumulation of LC-CoA under conditions of hyperlipidaemia and hyperglycaemia could contribute to the known insulin secretion defects under these conditions as K^+^ channel activation is expected to antagonise insulin secretion.

### Structural insights into the mechanism polyanionic lipid regulation

Based on crystallographic and functional data, is has been assumed that K_2P_ channels are regulated primarily via a highly dynamic SF gate located at the extracellular pore entrance (Bagriantsev et al., 2011; Piechotta et al., 2011; Bagriantsev et al., 2012; Brohawn et al., 2012; Miller and Long, 2012; Dong et al., 2015; Schewe et al., 2016; Lolicato et al., 2017; Schewe et al., 2019) but more recent structural studies revealed constriction sites in the ion permeations pathway below the SF in TASK-2 and TASK-1 channels (Li et al., 2020; Rodstrom et al., 2020). This raises the question which of the two possible gating structures is relevant in the context of the here reported polyanionic lipid regulation in these channels **(Figure 5A)**. In the TREK subfamily the PIP_2_/LC-CoA activation gate is likely the SF as lipid activation causes the transition of the voltage-dependent ion-flux gating mode of the SF into the leak mode displaying a linear I-V (**Figure 2B, Figure 5A, Supplementary Figure S1A-C, Supplementary Figure S2A,B**) (Schewe et al., 2016). However, THIK-1 channels display little voltage gating (unpublished results) and, thus, the principal role of the SF gate is unresolved and further experiments are required to determine the location of the PIP_2_/LC-CoA activation gate. Likewise, the structural mechanism of TRESK channel inhibition by LC-CoA requires further investigation.

For TASK-1 and TASK-3 channels our experiments suggest that inhibition by PIP_2_ and LC-CoA involves the lower X-gate crystallographically identified in TASK-1 channels (and assumed for TASK-3 channels), as the published mutations within this region also abolished PIP_2_ inhibition in both channels. However, the location of the PIP_2_/LC-CoA binding sites, as well as the possible overlap with the still unknown inhibitory DAG binding site, will require further investigation. In TASK-2 channels a lower gate mediating pH inhibition was recently identified in cryo-EM structures at a location similar to the X-gate in TASK-1 (Li et al., 2020). Although its involvement in PIP_2_/LC-CoA inhibition seems reasonable, mutation of a critical residue at the intracellular pH gate did not affect PIP_2_ inhibition (**Figure 4C, Supplementary Figure S1J**). Thus, additional studies are required to clarify the structural mechanisms of polyanionic lipid inhibition in TASK-2.

### Conclusions

This work establishes common polyanionic cellular lipids such as PIP_2_ and LC-CoA as regulators of channel activity for all 12 functionally expressing mammalian K_2P_ channels. We provide first insights into the location of the PIP_2_/LC-CoA inhibition gate in TASK-1 and TASK-3 but substantially more work is required to disclose the lipid binding sites in the various K_2P_ channels as well as the structural mechanism underlying the polyanionic lipid regulation. Finally, investigation of the relevant signal transduction pathways in native preparations is required to demonstrate the physiological relevance of linking K_2P_ channel activity to the complex metabolisms of phosphoinositides and fatty acids in the various tissues and cell types expressing K_2P_ channels.

## Materials and Methods

### Molecular biology and oocyte expression

For this study we used the coding sequences of hTWIK-1 (GenBank accession number: NM_002245.3), hTREK-1 (NM_172042.2), rTREK-1 (NM_172041.2), hTREK-2 (NM_138318.2), hTRAAK (AF_247042.1), hTALK-1 (NM_032115.4), hTALK-2 (EU978944.1), hTASK-2 (NM_003740.3), hTASK-1 (NM_002246.2), hTASK-3 (XM_011517102.1), hTHIK-1 (NM_022054), hTHIK-2 (NM_022055.1) and hTRESK (NM_181840.1). To increase surface expression and macroscopic currents, measurements of TWIK-1 and THIK-2 channels were done using channels with mutated retention motifs and a known activating mutation hTHIK-2, respectively (TWIK-1 I293A/I294A and THIK-2 R11A/R12A/R14A/R15A/R16A/A155P). All mutant channels were obtained by site-directed mutagenesis with custom oligonucleotides. All constructs were subcloned into the pFAW dual purpose vector suitable for *in vitro* transcription/oocyte expression and transfection of cultured cells and verified by sequencing. cRNA was synthetized using AmpliCap-Max T7 or SP6 High Yield Message Maker Kits (Cellscript, USA) and stored at −20°C (for frequent use) and −80 °C (for long term storage). *Xenopus laevis* oocytes were surgically removed from adult female frogs and treated with type II collagenase prior to manual defolliculation. Oocytes were injected with ~50 nl of channel-specific cRNA (0.5 - 1 μg*μl^−1^) and incubated at 17 °C for 1 - 14 days prior to the experimental use.

### Cell culture

HEK293 cells were cultured in Dulbecco’s Modified Eagles Medium (DMEM), supplemented with 10 % FCS and 10 U*ml^−1^ penicillin and 10 mg*ml^−1^ streptomycin in a 5 % CO2 atmosphere at 37 °C. The cells were transiently transfected with Lipofectamine 2000 (Invitrogen) in 24-well plates. For electrophysiological recordings the transfected cells were trypsinized at least 4 h before electrophysiological measurements and seeded onto sterile 10 mm coverslips in 35 mm culture dishes in antibiotic-free DMEM.

### Electrophysiology

Currents were recorded from inside-out membrane patches excised from cRNA-injected *Xenopus* oocytes or from transiently transfected HEK293 cells at room temperature (21 - 23 °C). For oocyte measurements, pipettes were made from thick-walled borosilicate glass capillaries and had resistances of 0.3 – 0.8 MΩ. Pipettes were filled with a standard pipette solution (in mM): 120 KCL, 10 HEPES and 3.6 CaCl_2_, pH 7.4 was adjusted with KOH/HCl. Currents were recorded using an EPC9 or EPC10 amplifier (HEKA electronics), sampled at 10 kHz and filtered with 3 kHz (−3 dB). The recording program was Patchmaster (HEKA electronics, version: v2×73.5). The used pulse protocols were either ramps ranging from −80 to +40 or +80 mV with a duration of 1 s and intervals of 4 or 9 s, or measurements were carried out using a continuous pulse at +40 mV. Solutions were applied to the cytoplasmic side of excised membrane patches via a gravity-flow multi-barrel application system. Standard intracellular (bath) solutions were composed of (in mM): 120 KCl, 10 HEPES, 2 EGTA and 1 Pyrophosphate, pH was adjusted to pH 7.4 with KOH/HCl. 5 mg*ml^−1^ BSA was added to obtain washout solution. Where indicated in experimental results, 120 mM KCl was replaced by 120 mM RbCl.

HEK293 cell measurements were done in the whole-cell configuration of the patch-clamp technique using an EPC10 amplifier (HEKA electronics) and the PatchMaster software (HEKA electronics, version: v2×73.5) with a sampling rate and filter as above. The cells were stimulated by a ramp protocol from −100 to +60 mV with 1 s duration and a 5 s interpulse duration. Pipette resistances ranged from 1 - 3 MΩ and pipettes were filled with intracellular solution (mM): 140 KCl, 2 MgCl_2_, 1 CaCl_2_, 2.5 EGTA, 10 HEPES, and pH 7.3 was adjusted with KOH/HCl. The bath contained (mM): 135 NaCl, 5 KCl, 2 MgCl_2_, 2 CaCl_2_, 10 glucose, 10 HEPES, and pH 7.3 was adjusted with NaOH/HCl. All modifying reagents were added directly to the bath to obtain the particular end concentrations. Lipids and other substances were stored as stock solutions (1 - 100 mM) at −20 °C and diluted in the bath solution to the final concentration prior to measurements.

### Chemicals, drugs and lipids

LC-CoA (14:0, 16:0, 18:0, 18:1, 18:2, 18:3, 22:0) were purchase from Avanti Polar Lipids (Alabaster, USA) and stock solutions were prepared in DMSO (1 - 5 mM). The phospholipase C activator m-3m3FBS, Tetrapentylammonium chloride (TPA) and L-α-phosphatidylinositol-4,5-bisphosphate ammonium salt (brain PI(4,5)P_2_; PIP_2_) were purchased from Sigma-Aldrich/Merck (Darmstadt, Germany) and stock solutions were prepared in DMSO (10 - 100 mM).

### Animals (*Xenopus laevis*)

The performed investigation conforms to the guide for the Care and Use of laboratory Animals (NIH Publication 85-23). For this study, we used female *Xenopus laevis* animals (25) that were accommodated at the animal breeding facility of the Christian-Albrechts-University of Kiel to isolate oocytes. Experiments using Xenopus toads are approved by the local ethics commission.

### Data analysis

Data analysis was done using Fitmaster (HEKA electronics, version: v2×73.5, Lamprecht, Germany), Microsoft Excel (Microsoft Corporation, version: Microsoft Office Professionel Plus 2019, Redmond, USA) and Igor Pro (Wavemetrics Inc., version: 6.3.7.2, Portland, USA).

Recorded currents were analysed from stable membrane patches at a voltage of +40 mV unless stated otherwise.

The fold activation of a ligand (drug or bioactive lipid) was calculated from the following equation:

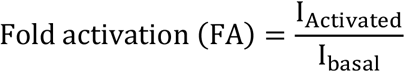

with I_Activated_ represents the stable current level in the presence of a given concentration of a respective ligand and I_basal_ the measured current before ligand application. Percentage inhibition of a ligand (drug or bioactive lipid) was calculated from stable currents of excised membrane patches using the following equation below:

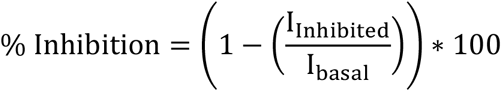

where I_Inhibited_ refers to the stable current level recorded in the presence of a given concentration of a drug or bioactive lipid and I_basal_ to the measured current before ligand application.

The macroscopic half-maximal concentration-inhibition relationship of a ligand was obtained using a modified Hill-equation depicted below:

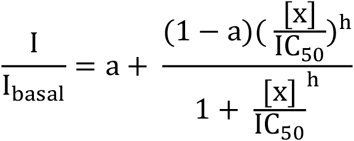

with I and I_basal_ are the currents in the absence and presence of a respective ligand, x is the concentration of the ligand, IC50 is the ligand concentration at which the inhibitory effect is half-maximal, h is the Hill coefficient and a is the fraction of unblockable current (a = 0 unless state otherwise). Data is represented throughout the manuscript as mean ± s.e.m. unless stated otherwise.

Image processing and figure design was done using the open source vector graphic program Inkscape (GNU General Public Licence, free Software Foundation, version: 1.0.1 (3bc2e813f5, 2020-09-07, https://inkscape.org, Boston, USA).

## Acknowledgement

These studies were supported by the Deutsche Forschungsgemeinschaft (DFG, German Research Foundation) to M.S. and T.B. as part of the Research Unit FOR2518, *Dynlon*.

## Author Contributions

M.S. and T.B. conceived and supervised the project; E.B.R., B.C.J., S.C. and M.S. performed patch-clamp experiments and analysed the data; M.M. created and supervised the generation of mutant channels; E.B.R. prepared all figures; E.B.R., M.S. and T.B. wrote the manuscript draft with critical comments of all authors.

## Open Research

The data supporting the findings of this study are available from the corresponding authors upon reasonable request.

## Supporting Information

**Supplementary Figure S1.**
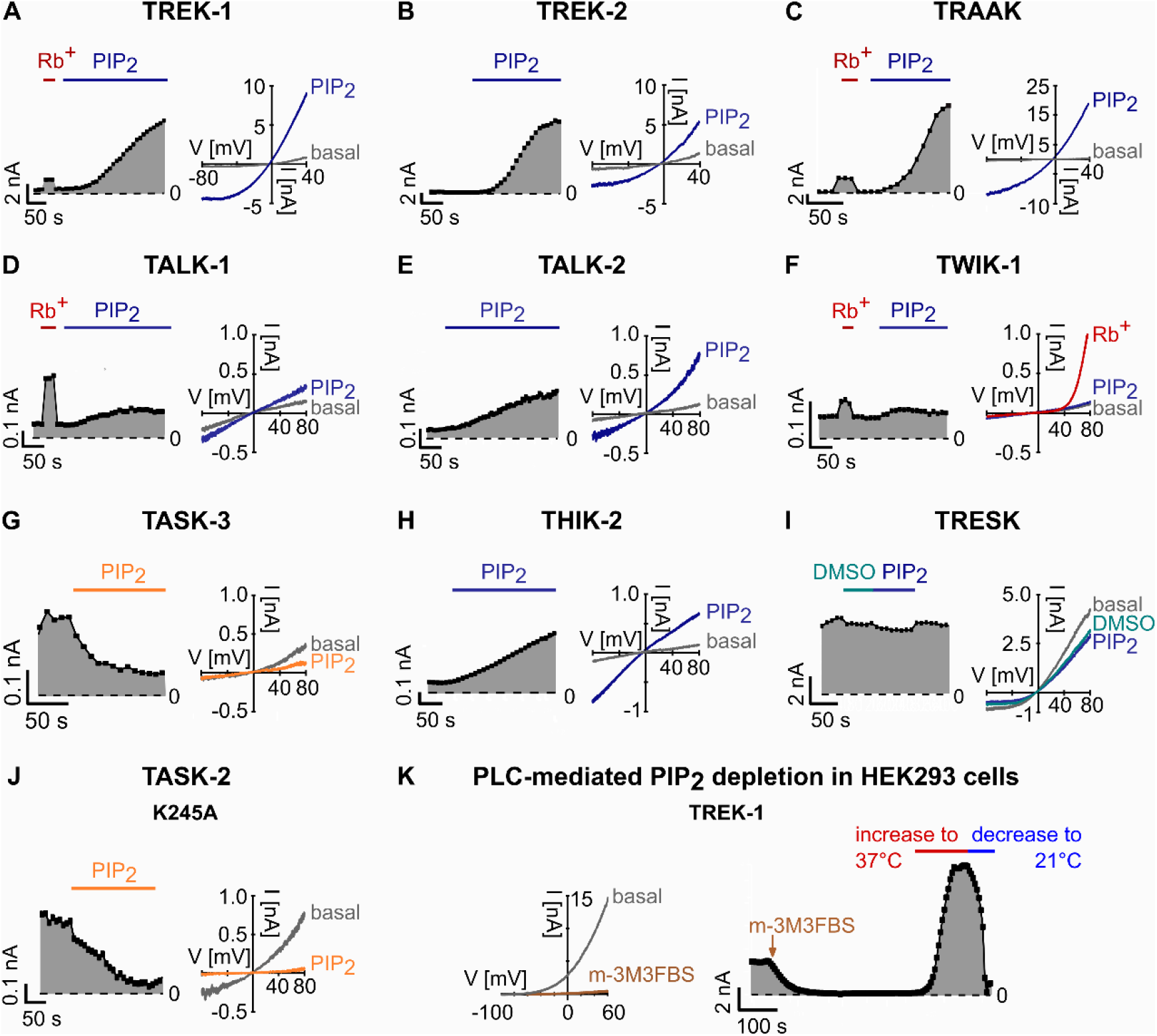
Representative traces of the PIP_2_ regulation of K_2P_ channels and PLC-mediated inhibition of TREK-1 channels. **A-J.** Representative current traces (right) and analysed currents at +40 mV plotted over time (left) of PIP_2_-activated (blue) TREK-1 (A), TREK-2 (B), TRAAK (C), TALK-1 (D), TALK-2 (E), THIK-2 and PIP_2_-inhibited (orange) TASK-3 (G) and TASK-2 K245A (J) K_2P_ channels. TWIK-1 (F) and TRESK (I) are not affected by PIP_2_. Currents were measured in voltage ramps between −80 and +80 mV in excised inside-out patches of *Xenopus* oocytes using K^+^ or Rb^+^ (red) bath solutions and in the presence of 10 μM PIP_2_ or DMSO (1 %) added to the standard K^+^ solution. **K.** Representative traces of TREK-1 currents in whole-cell experiments using HEK293 cells measured in voltage ramps between −100 and +60 mV and analysed at 0 mV. Measurements are performed in control solution and in the presence of 20 μM of the PLC activator m-3M3FBS. Additionally, the temperature was increased to 37 °C (red) and subsequently decreased (blue) to room temperature of 21 °C (blue).

**Supplementary Figure S2.**
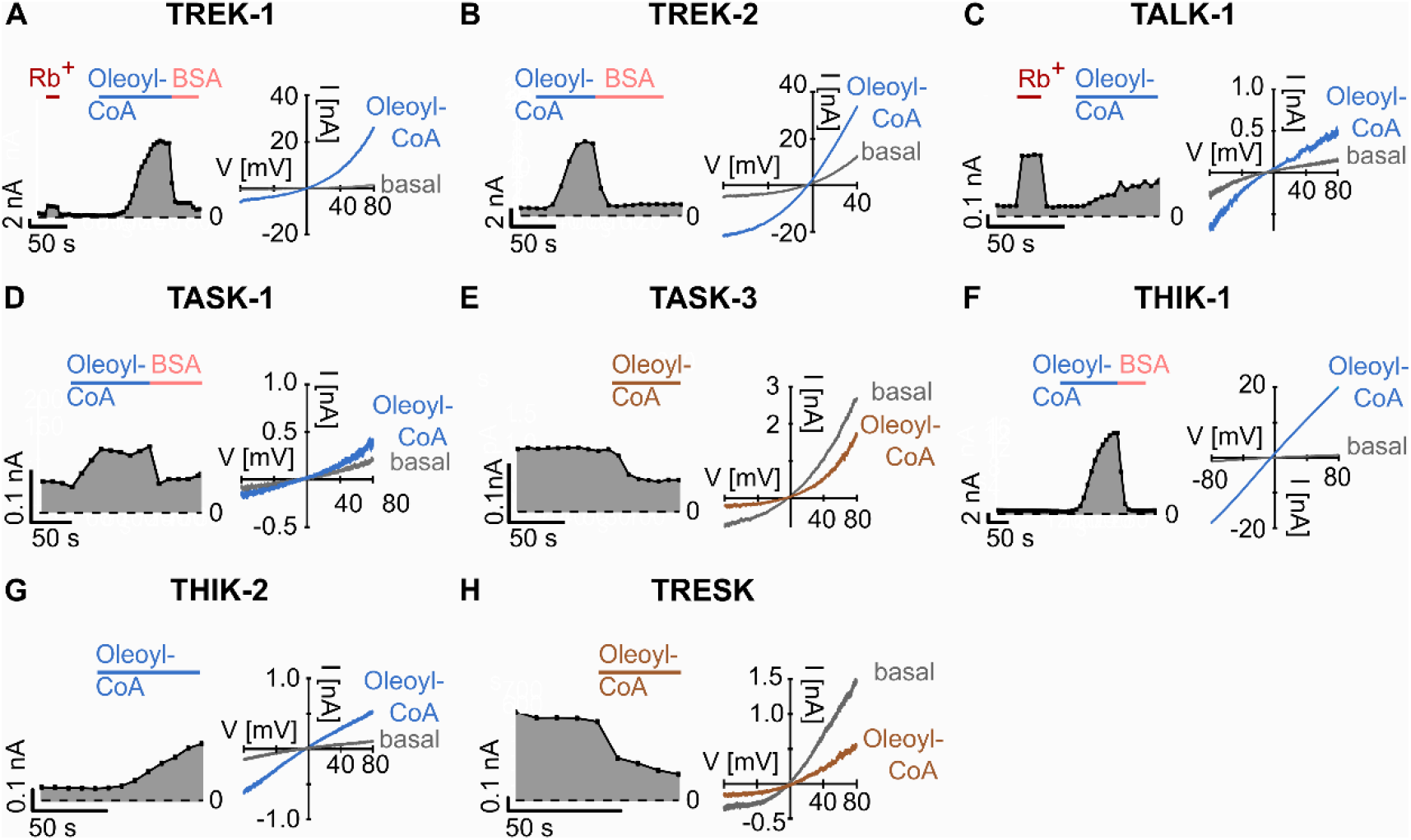
Oleoyl-CoA regulation of K_2P_ channels. **A-H.** Representative current traces (right) and analysed currents at +40 mV plotted over time (left) of oleoyl-CoA-activated (light blue) TREK-1 (A), TREK-2 (B), TALK-1 (C), TASK-1 (D), THIK-1 (F), and THIK-2 (G) and PIP_2_-inhibited (brown) TASK-3 (E) and TRESK (H) K_2P_ channels. Currents were measured in voltage ramps between −80 and +80 mV in excised inside-out patches of *Xenopus* oocytes using K^+^ or Rb^+^ (red) bath solutions and in the presence of 10 μM PIP_2_ or 5 mg*ml^−1^ BSA (pink) added to the standard K^+^ solution.

**Supplementary Figure S3.**
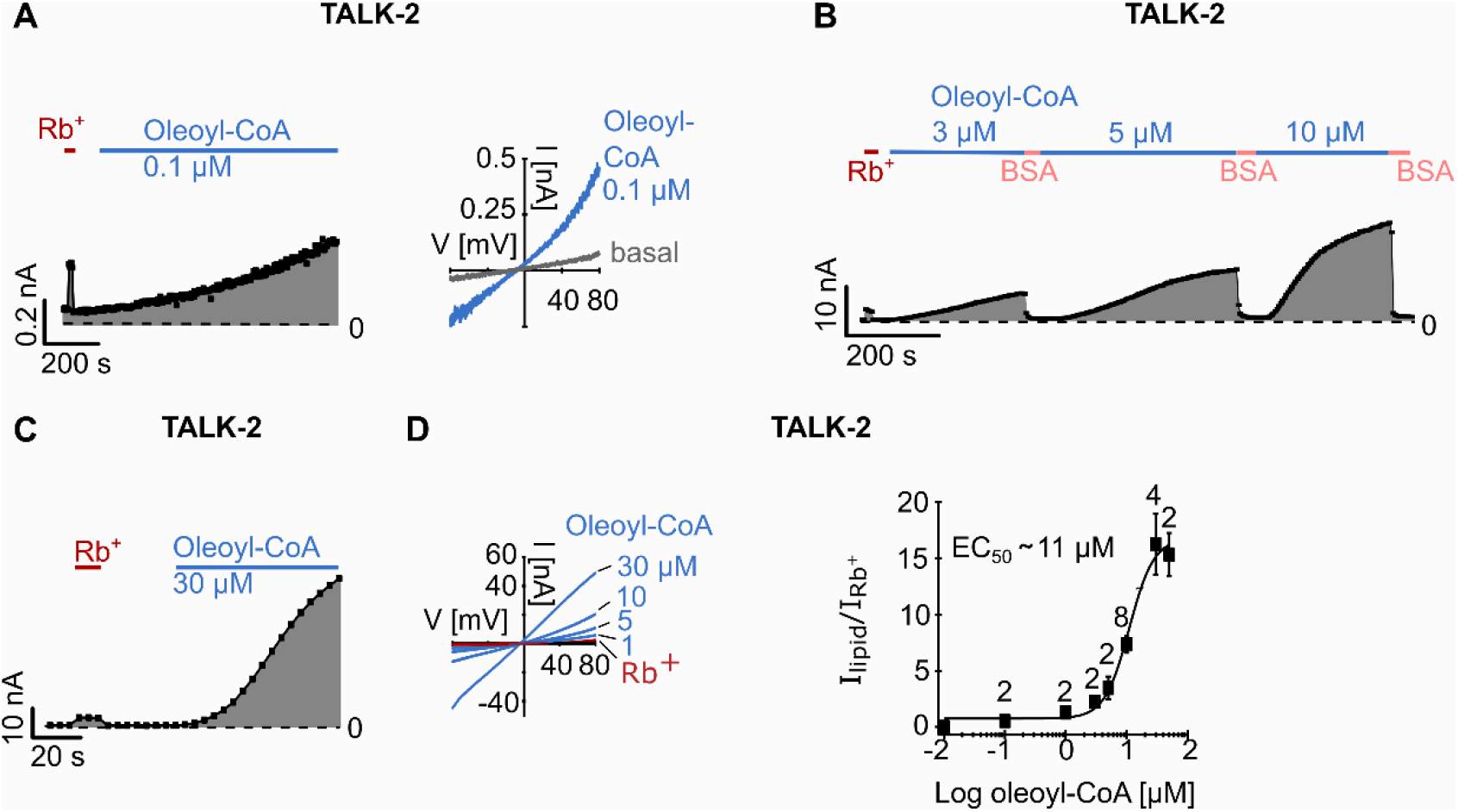
Oleoyl-CoA activation of TALK-2 K_2P_ channels. **A-C.** Representative current traces (right) and analysed currents at +80 mV plotted over time (left) of TALK-2 channels measured in voltage ramps between −80 and +80 mV in excised inside-out patches of *Xenopus* oocytes using K^+^ or Rb^+^ (red) bath solutions and in the presence of 0.1 μM (A), 3 μM, 5 μM and 10 μM (B) or 30 μM (C) oleoyl-CoA or 5 mg*ml^−1^ bovine serum albumin (BSA) for wash out. **D.** Dose-response relationship of oleoyl-CoA activation of TALK-2 channels measured as shown in the left panel (1, 5, 10, 30 μM oleoyl-CoA (blue), Rb^+^-activated (red) current). Note that all currents were normalised to the respective Rb^+^-activated current at +80 mV. The average fold activation by Rb^+^ (I_Rb_^+^/I_basal_) for TALK-2 WT channels is 12 ± 3 (n = 20). The EC50 of oleoyl-CoA was ~11 μM. Number (n) of independent experiments is indicated above the squares. All data is presented as mean ± s.e.m.

**Table S1.** Fold activation and % Inhibition induced by 10 μM PIP_2_ for the respective K_2P_ channel as depicted in Figure 1A.

| Fold activation |  |  |  |
| --- | --- | --- | --- |
| K<sub>2</sub>P channel | n | mean | s.e.m. |
| TREK-1 WT | 5 | 17.1 | 2.9 |
| TREK-2 WT | 5 | 10.9 | 4.0 |
| TRAAK WT | 7 | 113.6 | 24.4 |
| TALK-1 WT | 6 | 1.7 | 0.2 |
| TALK-2 WT | 15 | 4.4 | 0.9 |
| TASK-2 WT | 4 | 2.4 | 0.3 |
| THIK-1 WT | 5 | 19.3 | 2.3 |
| THIK-2 | 6 | 3.9 | 0.6 |
| TWIK-1 | 5 | 1.1 | 0.3 |
| Inhibition [%] |  |  |  |
| K<sub>2</sub>P channel | n | mean | s.e.m. |
| TASK-1 WT | 10 | 81.8 | 1.9 |
| TASK-2 WT | 11 | 82.8 | 2.8 |
| TASK-3 WT | 7 | 60.8 | 3.9 |
| TRESK WT | 6 | 6.9 | 1.1 |

**Table S2.**
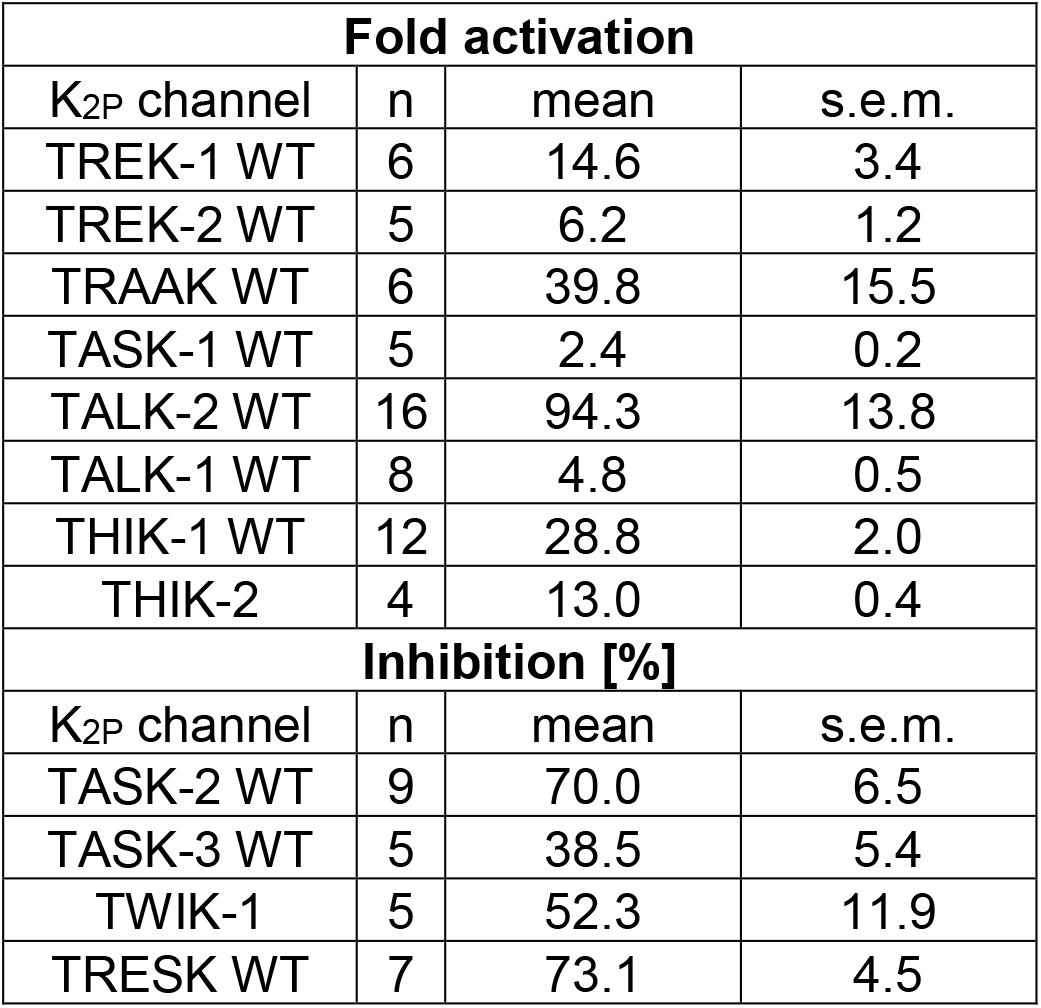
Fold activation and % Inhibition induced by 10 μM oleoyl-CoA for the respective K_2P_ channel as depicted in Figure 2A.

**Table S3.** Fold activation induced by the indicated LC-CoA species for TALK-2 K_2P_ channels as depicted in Figure 3D.

| **Fold activation by LC-CoA esters (3 μM) [$I_{lipid}/I_{Rb+}$]** |  |  |  |
| --- | --- | --- | --- |
| LC-CoA species | n | mean | s.e.m. |
| 18:3 | 3 | 0.3 | 0.1 |
| 18:2 | 3 | 0.7 | 0.2 |
| 18:1 | 3 | 4.4 | 0.8 |
| 18:0 | 3 | 11.7 | 2.0 |
| 14:0 | 4 | 0.1 | 0.04 |
| 16:0 | 3 | 3.0 | 0.7 |
| 18:0 | 3 | 11.7 | 2.0 |
| 22:0 | 3 | 7.8 | 1.8 |
| **Oleoyl-CoA dose response curve (normalised to Rb<sup>+</sup>)** |  |  |  |
| Concentration [μM] | n | mean [$I_{lipid}/I_{Rb+}$] | s.e.m. |
| 0.01 | 15 | 0.1 | 0.01 |
| 0.1 | 2 | 0.6 | 0.2 |
| 1 | 2 | 1.4 | 0.4 |
| 3 | 2 | 2.3 | 0.1 |
| 5 | 2 | 3.5 | 1.0 |
| 10 | 8 | 7.4 | 0.7 |
| 30 | 4 | 16.2 | 2.7 |
| 50 | 2 | 15.3 | 1.9 |

**Table S4.** Fold change induced by 10 μM PIP_2_ species for the respective K_2P_ channel as depicted in Figure 4D.

| **Fold change** |  |  |  |
| --- | --- | --- | --- |
| K<sub>2P</sub> channel | n | mean | s.e.m. |
| TASK-1 WT | 10 | 0.22 | 0.03 |
| TASK-1 L244A | 6 | 0.95 | 0.03 |
| TASK-1 R245A | 5 | 0.99 | 0.04 |
| TASK-3 WT | 7 | 0.39 | 0.04 |
| TASK-3 L244A | 5 | 1.22 | 0.08 |
| TASK-3 R245A | 5 | 1.36 | 0.05 |
| TASK-2 WT | 11 | 0.17 | 0.03 |
| TASK-2 K245A | 3 | 0.15 | 0.07 |

